# Action planning modulates the representation of object features in human fronto-parietal and occipital cortex

**DOI:** 10.1101/480574

**Authors:** Jena Velji-Ibrahim, J. Douglas Crawford, Luigi Cattaneo, Simona Monaco

**Affiliations:** CIMeC - Center for Mind/Brain Sciences, University of Trento, Via delle Regole 101, Trento, 38123, Italy; Center for Vision Research, York University, 4700 Keele Street, Toronto, Ontario, M3J 1P3, Canada; School of Kinesiology and Health Science, York University, 4700 Keele Street, Toronto, Ontario, M3J 1P3, Canada; Departments of Biology and Psychology, York University, 4700 Keele Street, Toronto, Ontario, M3J 1P3, Canada

**Keywords:** action planning, object features, fronto-parietal cortex, early visual cortex, functional magnetic resonance imaging (fMRI), multivoxel pattern analysis, decoding

## Abstract

The role of the visual cortex has been extensively studied to determine its role in object recognition but to a lesser degree to determine how action planning influences the representation of objects’ features. We used functional MRI and pattern classification methods to determine if during action planning, object features (orientation and location) could be decoded in an action-dependent way. Sixteen human participants used their right dominant hand to perform movements (Align or Open reach) towards one of two 3D-real oriented objects that were simultaneously presented and placed on either side of a fixation cross. While both movements required aiming toward target location, Align but not Open reach movements required participants to precisely adjust hand orientation. Therefore, we hypothesized that if the representation of object features is modulated by the upcoming action, pre-movement activity pattern would allow more accurate dissociation between object features in Align than Open reach tasks. We found such dissociation in the anterior and posterior parietal cortex, as well as in the dorsal premotor cortex, suggesting that visuomotor processing is modulated by the upcoming task. The early visual cortex showed significant decoding accuracy for the dissociation between object features in the Align but not Open reach task. However, there was no significant difference between the decoding accuracy in the two tasks. These results demonstrate that movement-specific preparatory signals modulate object representation in the frontal and parietal cortex, and to a lesser extent in the early visual cortex, likely through feedback functional connections.

## Introduction

To execute actions in daily life successfully, our brain needs to obtain accurate information about the orientation, location, shape and size of a target object. Picking up a pen, for example, would be more successful when one is focused on its orientation rather than its color. Considerable research has investigated the role of fronto-parietal reaching and grasping networks in successfully executing actions (for reviews see: Vesia and Crawford 2012; Gallivan and Culham 2015), and multivoxel pattern analysis has allowed examining the representation of action intention in fronto-parietal and temporal-occipital cortices seconds before participants start to move (Gallivan et al. 2011; Gallivan, Chapman, et al. 2013; Monaco et al. 2019). Action planning strongly relies on the representation of our surrounding for generating accurate and effective movements, and at the same time, it enhances the detection of features that are relevant for behaviour (Gutteling et al. 2011). This suggests that feedback connections from frontal and parietal areas involved in action preparation modulate the activity in visual areas (Gutteling et al. 2013), mediating the enhancement of feature perception during action planning and allowing action-relevant information to be shared between visual and somato-motor areas. The outstanding question is how early in the visual system, in terms of cortical location, is this modulation detected.

Human neuroimaging studies have shown that the early visual cortex (EVC) is reactivated at the time of delayed actions despite the absence of visual information (Singhal et al. 2013; Monaco et al. 2017). The re-recruitment of the EVC during action execution might enhance the processing and retrieval of object features for subsequent object manipulations. Another possible explanation of these results is that the somatosensory and motor feedback elicited during the execution of a movement might elicit detectable responses in the visual cortex despite the absence of online visual information. However, evidence from electrophysiology has shown preparatory activity in visual areas shortly before action execution (van Elk et al. 2010), when somatosensory feedback is not available yet. In addition, fMRI studies show that the activity pattern in the EVC allows decoding different action plans towards the same object (Gutteling et al. 2015; Gallivan et al. 2019) and that these results cannot be merely explained by imagery (Monaco et al. 2020). Overall, these findings indicate that the EVC might be involved in more than just low-level feature processing for action planning.

The goal of this study was to determine whether the representation of object features, such as orientation, varies as a function of the planned action. And if so, whether this is the case only in sensorimotor areas of the fronto-parietal network, or even in the occipito-temporal and early visual cortex. To test this, we used a paradigm in which participants performed one of two actions towards one of two objects that differed in location and orientation and were concurrently presented. While object location was relevant for both motor tasks, object orientation was relevant only for one of the two tasks. To uncover influences of action preparation on processing of object features we used multi-voxel pattern analysis (MVPA) on the planning phase preceding the action. We hypothesized that during movement preparation areas that play a specific role in processing action relevant features of objects (i.e., orientation) would be modulated in a task-specific fashion.

## Methods

Our main question was aimed to investigate whether action planning modulated the activity pattern elicited by the visual presentation of two simultaneously presented objects. If the representation of an object is shaped by the intended action, we would see enhanced dissociation between the two oriented objects when participants were planning an action that had to be adjusted to the orientation of the object (Align) as compared to a movement for which object orientation was irrelevant (Open reach) (Figure 1, left panel). Therefore, we examined the activity pattern in areas of ventral and dorsal visual stream known to have a representation of action planning. Further, to explore whether the modulation might occur as early as in the EVC, we examined the activity pattern in the Calcarine sulcus, corresponding to the peripheral location of the objects in the visual field, as well as in the Foveal cortex, corresponding to central vision. In fact, previous studies have shown that information about the category of objects viewed in the periphery is fed back to the foveal retinotopic cortex and correlates with behavioral performance (Williams et al. 2008; Fan et al. 2016). Therefore, we tested whether action-relevant features of objects presented in the periphery are also distinguishable in the foveal cortex by identifying the corresponding retinotopic locations using retinotopic mapping procedures.

**Figure 1.**
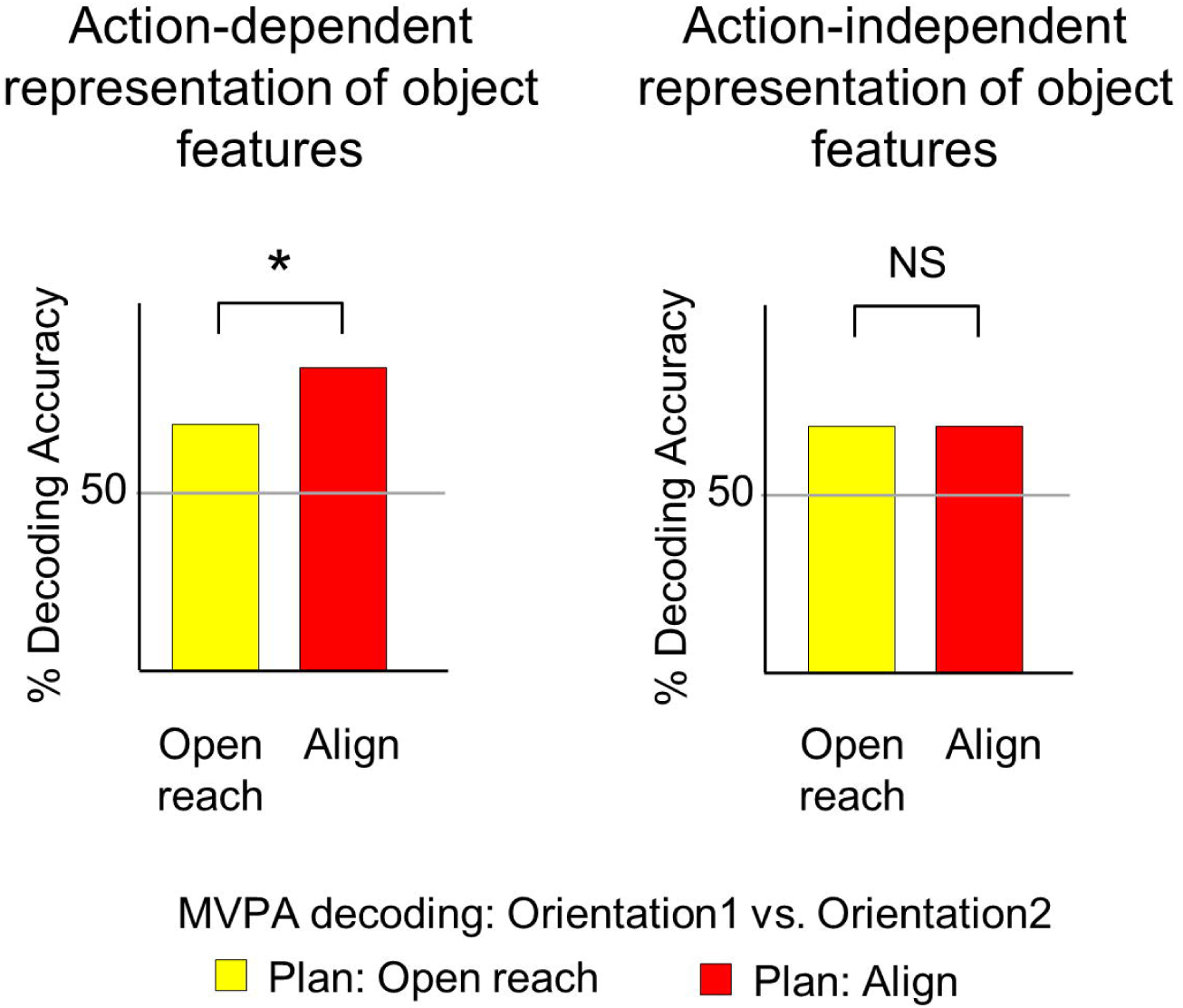
Hypothesis. Predicted percent decoding accuracies based on MVPA for the dissociation between two differently oriented objects in Align and Open reach movements during action planning. We hypothesized that action-dependent modulation of the representation of object features would be reflected in higher decoding accuracies for Align then Open reach tasks (left panel).

### Sessions

The experimental and retinotopic mapping sessions took place on two different days. The experimental session lasted approximately 2 hours, including screening and set-up time, while the retinotopic mapping session took approximately 30 minutes to be completed.

### Participants

Twenty-six right-handed volunteers (14 females) participated in this study. The age range of participants was 20-45, with an average age of 30.4 years. Sixteen participants volunteered for the experimental runs and 14 participants took part in the independent localizer runs for retinotopic mapping. Four of these participants took part in both sessions. All participants had normal or corrected-to-normal vision and none of the participants had any known neurological deficit. All participants provided informed consent and approval was obtained from the ethics committee for experiments involving human participants at the University of Trento.

### Experimental Setup: Apparatus and Stimuli

The experimental set up is illustrated in Figure 2A. Participants lay on the bed of a 4-Tesla MRI scanner and performed actions towards one of two real 3D objects. Both objects were affixed to strips of Velcro attached to a platform that was covered with the complementary side of Velcro. The platform was placed over the pelvis of the participant. This device enabled subjects to perform hand actions (Align and Open reach movements) towards two wooden objects mounted on the platform. The two objects were placed on either side of a fixation cross. The object on the left was oriented at about -45° and will be referred to as counterclockwise-left (CCW-left) while the one on the right was oriented approximately at 45° and will be referred to as clockwise-right (CW-right) (Figure 2B). The head of the participant was slightly tilted (∼30°) to allow direct viewing of the stimuli presented on the platform. The platform was perpendicular to gaze. The fixation cross was placed ∼7.5 cm above the object at a viewing distance of ∼65 cm, such that the objects were at eccentricities greater than 6.6 degrees of visual angle on both sides of fixation. The platform covered the entire portion of the lower visual field. The fixation point was located just below the bore of the magnet, such that the bore was on the upper visual field. Therefore, the visual field was almost entirely covered by the platform (lower part) and the bore of the magnet (upper part).

**Figure 2.**
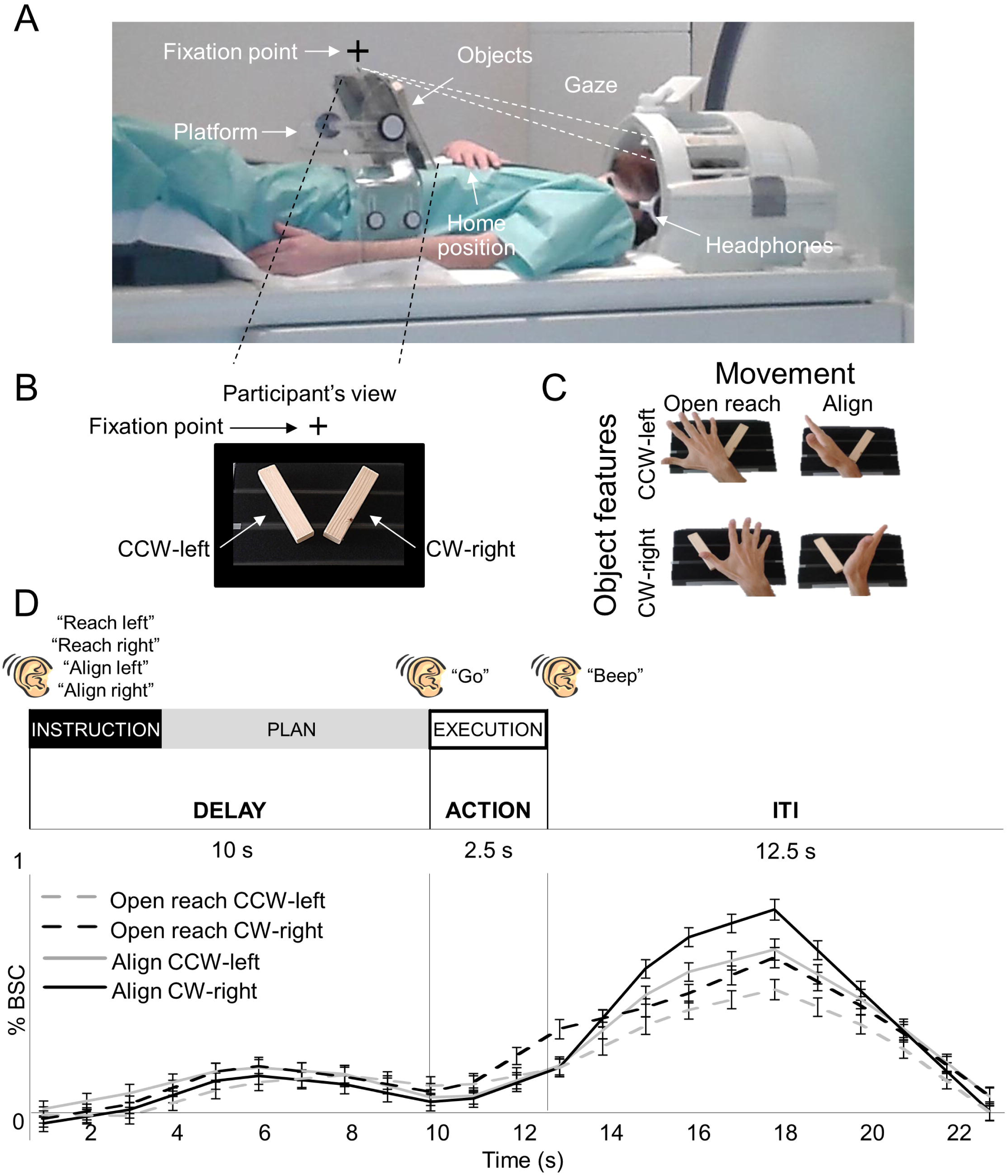
Image of the experimental setup, design, paradigm, and timing. (A) The setup required participants to gaze at the fixation point, marked with a cross, while preforming the task. (B) Participant’s view of the platform with the two oriented objects: counterclockwise-left (CCW-left) and clockwise-right (CW-right). (C) Experimental design. Participants performed four actions towards the instructed oriented object. Movements consisted of Align or Reach towards the CCW- left or CW-right object. As shown here, Align required the precise adjusting of a participant’s hand over the object while Reach movements were coarse. (D) Experimental paradigm and timing. Each trial consisted of three phases: instruction, plan and execution. At the beginning of each trial, an auditory cue indicated the condition type to the participant (“Reach Left”, “Reach Right”, “Align Left”, and “Align Right”,). There was a delay of 10 seconds during which participants did not perform any action until they heard a “go” cue upon which they performed the movement that they had been instructed at the beginning of the same trial. The end of the trial was cued by a “beep” sound, which prompted participants to return the hand to the home position. We used a 12.5 second intertrial interval. We focused our analysis on the 7.5 sec preceding action execution, during the plan phase. Lower panel: Group-averaged % BOLD signal change extracted from the calcarine sulcus in the left hemisphere for Align and Open reach CCW-left and CW-right. Error bars indicate standard errors.

To limit motion artifacts, the right upper arm was supported with foam and gently restrained. Reaches were thus performed by movements of the right forearm and hand. A button box was placed around the participants’ abdomen and served as the starting point for each trial. Hand actions were monitored with a Sony HDR-UX1E digital video camera. The lights were on throughout the experiment and the hand was visible to participants. Participants wore headphones to hear auditory instructions and cues.

### Experimental Paradigm

We used functional magnetic resonance imaging (fMRI) to measure the blood-oxygenation-level dependent (BOLD) signal (Ogawa et al. 1992) in a slow event-related delayed-action paradigm. We had a 2 x 2 factorial design, with factors of Movement (Align or Open reach) and Object features, such as orientation and location (CCW-left or CW-right), which lead to four conditions: Align CCW-left, Align CW-right, Open reach CCW-left and Open reach CW-right (Figure 2C). As shown in Figure 2D, each trial began with an auditory instruction indicating to the participant the action type and the object to be acted upon at the end of the trial. The auditory instructions were: Align left, Align right, Reach left, and Reach right. Then there was a delay of 10 seconds during which participants did not perform any action until they heard a go cue. When hearing the go cue, participants had 2.5 seconds to perform the movement that they had been instructed at the beginning of the same trial. A beep sound cued the participant to return the hand to the home position on the button box where it rested until the next trial began. The inter trial interval (ITI) lasted 12.5 seconds. The delayed timing of the experiment allowed us to isolate the pre-movement activity during the planning phase before the execution of the movement.

Participants were instructed to fixate their eyes on the fixation cross throughout the experiment. The objects were always visible to the participants throughout the trial. At the beginning of each run there were 17.5 s (7 volumes) during which participants rested. This phase was include in the baseline, together with the ITI. Align movements consisted of reaching to the CCW-left or CW-right object and adjusting the hand precisely over it, as in a manual orientation matching task. The Open reach movements consisted of moving the hand above the instructed object with an open palm. Therefore, while both movements were directed to one of the two object locations, only Align movements also required adjusting the hand according to the orientation of the object. In both movement types, participants touched the object during the execution of the movement. Each participant was trained and tested in a short practice session (10-15 minutes) prior to the fMRI experiment. The hand was monitored with a camera to confirm that participants were performing the correct tasks during the fMRI experiment.

Each run included 7 trials per experimental condition, for a total of 28 trials per run. Each trial type was presented in counterbalanced order for a run time of ∼12.5 minutes. Participants completed 5 functional runs for a total of 140 trials per subject (35 trials per condition).

### Imaging Parameters

This study was conducted at the University of Trento’s Center for Mind/Brain Sciences (CIMeC) in Mattarello, Italy using a 4T Bruker MedSpec whole body MRI system (Bruker BioSpin, Ettlingen, Germany), equipped with Siemens Magnetom Sonata gradients (200 T/m/s slew rate, 40 mT/m maximum strength; Siemens Medical Solutions, Erlangen, Germany) and an eight-channel head coil. Functional data was acquired using T2*-weighted segmented gradient echo-planar imaging sequence (repetition time [TR] = 2500 ms; echo time [TE] = 33 ms; flip angle [FA] = 78° for experimental runs, 73° for eccentricity mapping; field of view [FOV] = 192 × 192 mm, matrix size = 64 × 64 leading to an in-slice resolution of 3 × 3 mm; slice thickness = 3 mm, .45mm gap). Each volume was comprised of 35 slices for experimental runs, 33 slices for eccentricity mapping which were collected in interleaved order. During each experimental session, a T1-weighted anatomical reference volume was acquired using a MPRAGE sequence (TR = 2700 ms; TE = 7°; inversion time TI = 1,020 ms; FA = 7°; FOV = 256 x 224 x 176, 1 mm isotropic resolution).

### Preprocessing

Data was analyzed using Brain Voyager QX software version 2.8.4 (Brain Innovation, Maastricht, Netherlands). The first 3 volumes of each functional scan were discarded to avoid T1 saturation effects. For each run, slice scan time correction (cubic spline), temporal filtering (removing frequencies <2 cycles/run), and 3D motion correction (trilinear/sinc) were performed. To complete 3D motion correction, each volume of a run was aligned to the volume of the functional scan that was closest in time to the anatomical scan. Seven runs showing abrupt head movements greater than 1mm were discarded from the analyses. The data was transformed into Talairach space (Talairach and Tournoux 1988).

### General Linear Model (GLM)

We analyzed the data from the experimental runs using a group random-effects (RFX) general linear model (GLM) that included one predictor for each of the four conditions (Align CCW-left, Align CW-right, Open reach CCW-left, Open reach CW-right) and three trial phases (Instruction, Plan, and Execution) resulting in a total of 12 predictors of interest. In addition, we included 6 motion correction parameters as predictors of no interest. Each predictor was derived from a rectangular-wave function (1 volume or 2.5 s for the Instruction phase, 3 volumes or 7.5 s for the Plan phase phase and 1 volume or 2.5 s for the Execution phase) convolved with a standard hemodynamic response function (HRF; Brain Voyager QX’s default double-gamma HRF). The GLM was performed on %-transformed beta weights (β).

### Regions of Interest (ROIs)

We localized nine ROIs in the left and right hemisphere of each participant to determine whether action-relevant features of objects (orientation and location) can be distinguished from the activity pattern elicited by action planning, before participants performed the action (see “Localization of ROIs” for details about the procedure of localization). We identified five areas in ventral and dorsal visual stream that are typically involved in visually guided actions: superior-parietal occipital cortex (SPOC), anterior intrapariental sulcus (aIPS), posterior intraparietal sulcus (pIPS), lateral occipital cortex (LOC) and dorsal premotor cortex (dPM). We chose these ROIs based on their involvement in: 1) adjusting hand orientation during action execution in humans (SPOC, dPM, pIPS: (Monaco et al. 2011)) and macaques (aIPS: (Baumann et al. 2009); V6A/SPOC: (Battaglini et al. 2002; Fattori et al. 2009); 2) processing grasp-relevant dimensions of objects (SPOC, dPM and LOC: (Monaco et al. 2014)); and 3) discriminating object orientation (LOC: (Ganel and Goodale 2017)). Further, we localized three areas in the EVC: the Calcarine sulcus, Foveal cortex and the dorsal secondary visual cortex (V2d). Recent fMRI studies have found that actions intention can be decoded from the retinotopic location of the target object in the EVC when the object is located in the peripheral visual field, corresponding to the Calcarine sulcus (Gallivan et al. 2019; Monaco et al. 2020). Therefore, we localized the retinotopic location of the objects along the calcarine sulcus to explore whether there is an action-dependent representation of object features. Further, behavioural and transcranial magnetic stimulation studies have shown that information about objects located in the visual periphery is fed back to the foveal retinotopic cortex, corresponding to central vision, and correlates with behavioral performance (Williams et al. 2008; Fan et al. 2016). Therefore, we localized the part of retinotopic cortex corresponding to central vision in the Foveal cortex. In addition, we localized V2d to examine the representations in other areas of the EVC, as well as area MT, which is sensitive to the motion of visual stimuli (Zeki et al. 1991). Since we could not track eye-movements during the experiment, significant decoding accuracy in MT during action planning might suggest that participants performed saccades despite the instruction to fixate the fixation cross.

### Multi-Voxel Pattern Analysis (MVPA)

#### Linear Discriminant Analysis (LDA) single trial classification

MVPA was performed to determine if actions modulate the activity pattern in our ROIs through the decoding of object features during the planning phase preceding the two actions. In particular, in areas that showed movement-dependent representation of object features during action planning, we expected significant decoding accuracy for the dissociation of CCW-left vs. CW-right in Align but not Open reach conditions. Importantly, we expected a higher decoding accuracy for the dissociation of object features for Align than Open reach conditions (Figure 1, left panel). On the other hand, in areas that show movement-independent representation of object features, we would expect no significant difference in the dissociation of objects features for Align and Open reach conditions (Figure 1, right panel).

We used a combination of in-house software (using MATLAB) and the CoSMo MVPA Toolbox for MATLAB (http://cosmomvpa.org), with an LDA classifier (http://cosmomvpa.org/matlab/cosmo_classify_lda.html#cosmo-classify-lda). For each participant, we estimated a GLM on unsmoothed data modeling every trial per condition. The 4 experimental conditions (Align CCW-left, Align CW-right, Open reach CCW-left, Open reach CW-right) by 3 phases of the trial (Instruction, Delay, Action) by 7 repetitions per run by 5 runs, gave rise to a total of 420 regressors of interest per subject. In addition, we modelled movement parameters (3 rotations and 3 translations) as predictors of no interest. We adopted a ‘leave-one-run-out’ cross-validation approach to estimate the accuracy of the LDA classifier.

### Classifier inputs

To provide inputs for the LDA classifier, the β weights were extracted from the phase of interest (i.e. Plan or Execution phase) for each voxel in the ROI. Each phase included the volumes defined in the predictors for the GLM estimated on unsmoothed data. In particular, the Plan phase consisted of 3 volumes following the Instruction phase, while the Execution phase consisted of 1 volume following the Plan phase.

### Cross-decoding

We examined whether object information is encoded in similar ways in the two Action conditions (Align and Open reach) by testing whether the LDA classifier trained to discriminate between two objects (CCW-Left vs. CW-Right) in one of the two Action types could then be used to accurately predict trial identity when tested on the other Action type. The trials of the train and test runs were taken from different Action conditions such that the training was performed considering the pairwise comparison between the two objects (CCW-Left vs. CW-Right) in the Open reach condition and tested in the Align condition, and vice versa.

### Statistical analysis

We statistically assessed decoding significance across participants with a two-tailed t-test versus 50% chance decoding. To further explore whether the decoding accuracy was higher for the dissociation between the two objects in Align than Open reach movements, we performed two-tailed paired sample t-tests between the decoding accuracies in the two movement types. To control for multiple comparisons and reduce Type I errors, a false discovery rate (FDR) correction of q ≤ .05 was applied, based on the number of ROIs and the number of t-tests performed within each time phase (Benjamini and Yekutieli 2001). We report the results that did not survive FDR correction, but only discuss FDR-corrected results.

We used G*Power to perform a post-hoc power analysis based on a recent study that has shown significant decoding accuracy for the dissociation between different action plans in the EVC (Monaco et al. 2020). We used the effect size (Cohen’s d 0.83) from their left EVC ROI and calculated that two-tailed t-tests with 16 participants would provide a power of 0.87.

### Voxelwise Analysis

Since our ROIs are known to be involved in visually guided actions, we localized them with a contrast of: (Action execution > baseline). Activation maps for group voxelwise results were overlaid on the average inflated brains of all participants by cortex-based alignment (Figure 3A).

**Figure 3.**
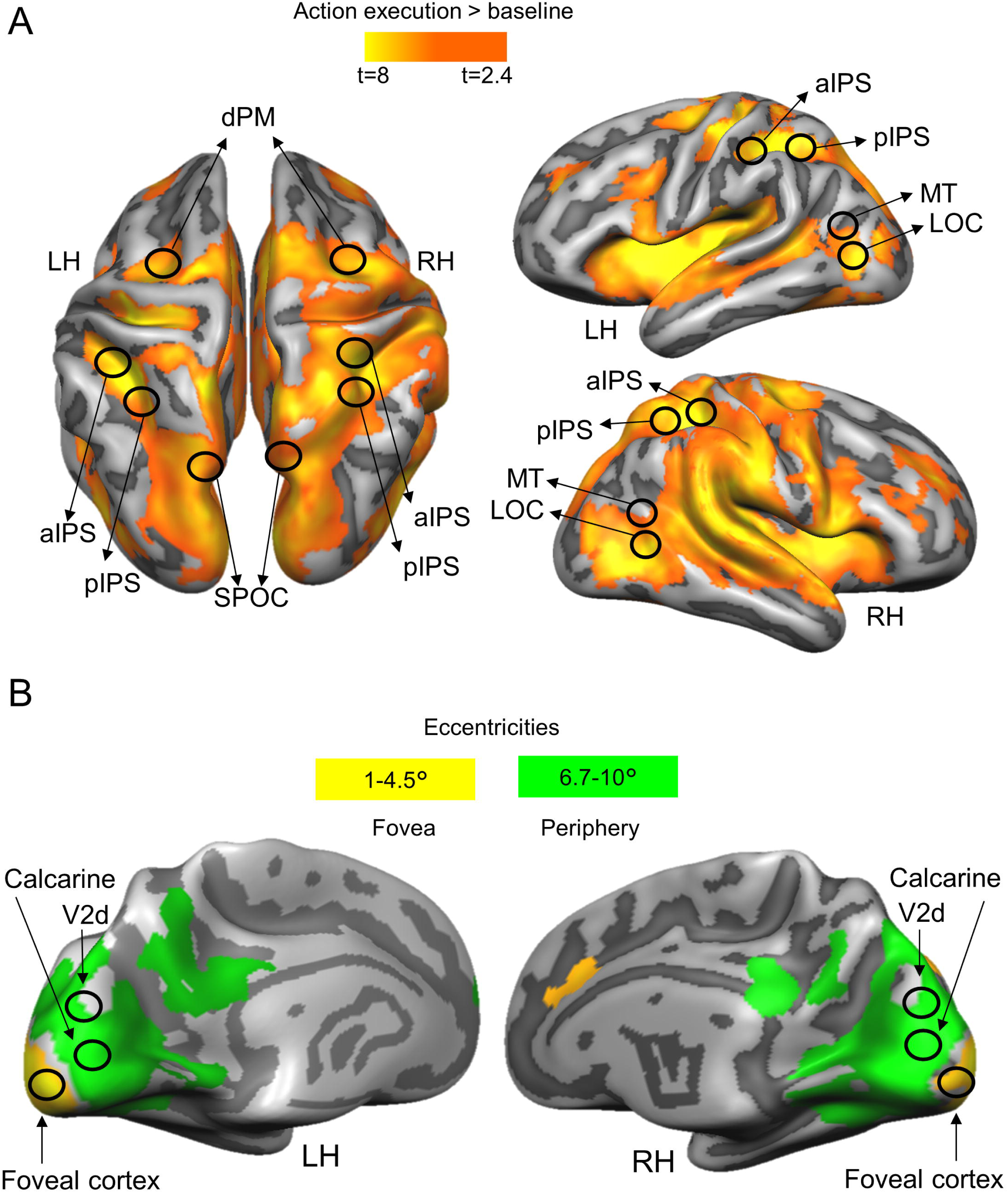
Activation maps overlaid on the average cortical surface. (A) Activation maps for the localization of ROIs in dorsal and ventral stream obtained with the univariate contrast: Action execution > baseline. Voxelwise statistical maps were obtained with the Random Effects GLM of experimental runs. (B) EVC activation maps obtained with the eccentricity mapping runs. Areas with higher activation for 6.7-10° than 1-4.5° (green) and areas with higher activation for eccentricities 1-4.5° than 6.7-10° (yellow). Eccentricity mapping was used to independently localize the area slightly above the Calcarine sulcus (which corresponds to the objects’ placement in the visual field) and the Foveal cortex (which corresponds to central vision). Eccentricity mapping was completed separately from the experiment on a set of 14 participants.

To correct for multiple comparisons, we performed cluster threshold correction for each activation map generated with a voxelwise contrast by using Brain Voyager’s cluster-level statistical threshold estimator plug-in (Forman et al. 1995; Goebel et al. 2006). This algorithm applied 1000 iterations of Monte Carlo simulations to estimate the number of neighboring false positive voxels which were active purely due to chance. Areas that did not survive this correction were excluded from the analyses.

### Retinotopic Mapping

In a separate session, a set of 14 participants underwent eccentricity mapping procedures. Of these participants, 4 also took part in the experiment. The expanding ring, used for eccentricity mapping, increased logarithmically as a function of time in both size and rate of expansion, so as to match the estimated human cortical magnification function (for details see (Swisher et al. 2007)). The smallest and largest ring size corresponded, respectively, to 1° and 10° of diameter. We divided the 10° into 8 equal time bins (of 8 seconds each). The eccentricity mapping localizer was composed of 8 cycles, each lasting 64 seconds. A fixation time was added at the beginning and at the end of the experiment for a total duration of 9 minutes per run. The stimuli were rear-projected with an LCD projector (EPSON EMP 7900 projector; resolution, 1280x1024, 60-Hz refresh rate) onto a screen mounted behind the participants’ head. The participants viewed the images through a mirror mounted to the head coil directly above the eyes. For eccentricity stimuli, we convolved a boxcar-shaped predictor for each bin with a standard HRF and performed contrasts using an RFX GLM.

We present results of eccentricity mapping because our hypotheses are in terms of the eccentric locations rather than the specific visual areas implicated. Moreover, the Occipital pole corresponds to the foveal confluence of several retinotopic visual areas, specifically V1, V2, and V3 (Wandell et al. 2007; Schira et al. 2009).

### Localization of ROIs

For each participant, we localized individual ROIs in two main steps. In the first step, we outlined the areas based on the group activation map obtained with the RFX-GLM contrast: (Action execution > baseline), by circumscribing group ROIs (9-mm radius) around their expected anatomical landmarks (Figure 3). Dorsal and ventral stream ROIs were localized at the: superior end of the parietal occipital sulcus for SPOC; junction of intraparietal and postcentral sulci for aIPS; posterior end of the intraparietal sulcus for pIPS; T-junction of superior frontal and precentral sulci for dPM; junction of inferior temporal sulcus and lateral occipital sulcus for LOC, and at the intersection of the occipital, temporal and parietal lobe for MT (Zeki et al. 1991) (Figure 3A). The group EVC ROIs for the Calcarine sulcus and Foveal cortex were selected based on the overlap between the activation map obtained with the contrast (Action execution > baseline) and the one resulting from the eccentricity mapping. We reasoned that since the object was located at ∼6.6° of visual angle below the fixation point, we could localize its location in the visual cortex at eccentricities greater than 6.7° of visual angle on or slightly above the Calcarine sulcus, consistent with the location of the object in the lower visual field (Figure 3B, green activation map). To localize the Foveal cortex corresponding to central vision, we selected voxels that showed higher activation for eccentricities up to 4.5° than greater eccentricities (Wandell 1995; Strasburger et al. 2011) (Figure 3B, yellow activation map). In order to determine the portion of the EVC corresponding to V2d, we used a published probabilistic atlas (Wang et al. 2015) that provides a dataset with the full probability maps of topographically organized regions in the human visual system (www.princeton.edu/~napl/vtpm.htm). In particular, the atlas provides the probabilistic maps generated from a large population of individual subjects (N = 53) tested with standard retinotopic mapping procedures and allows defining the likelihood of a given coordinate being associated with a given functional region for results obtained from any independent dataset once transformed into the same standard space. Therefore, we converted the probabilistic maps from MNI to Talairach space and used them to define V2d. In the second step, within each group ROI we defined individual ROIs, separately for each participant, as spheres with radius of 6 mm centered around each individual peak voxel resulting from the single-subject GLM contrasts (Action execution > baseline). This approach ensured that all regions were objectively selected, and that all ROIs had the same number of anatomical voxels (925 mm^3^). The averaged Talairach coordinates of individual ROIs are shown in Table 1.

**Table 1.**
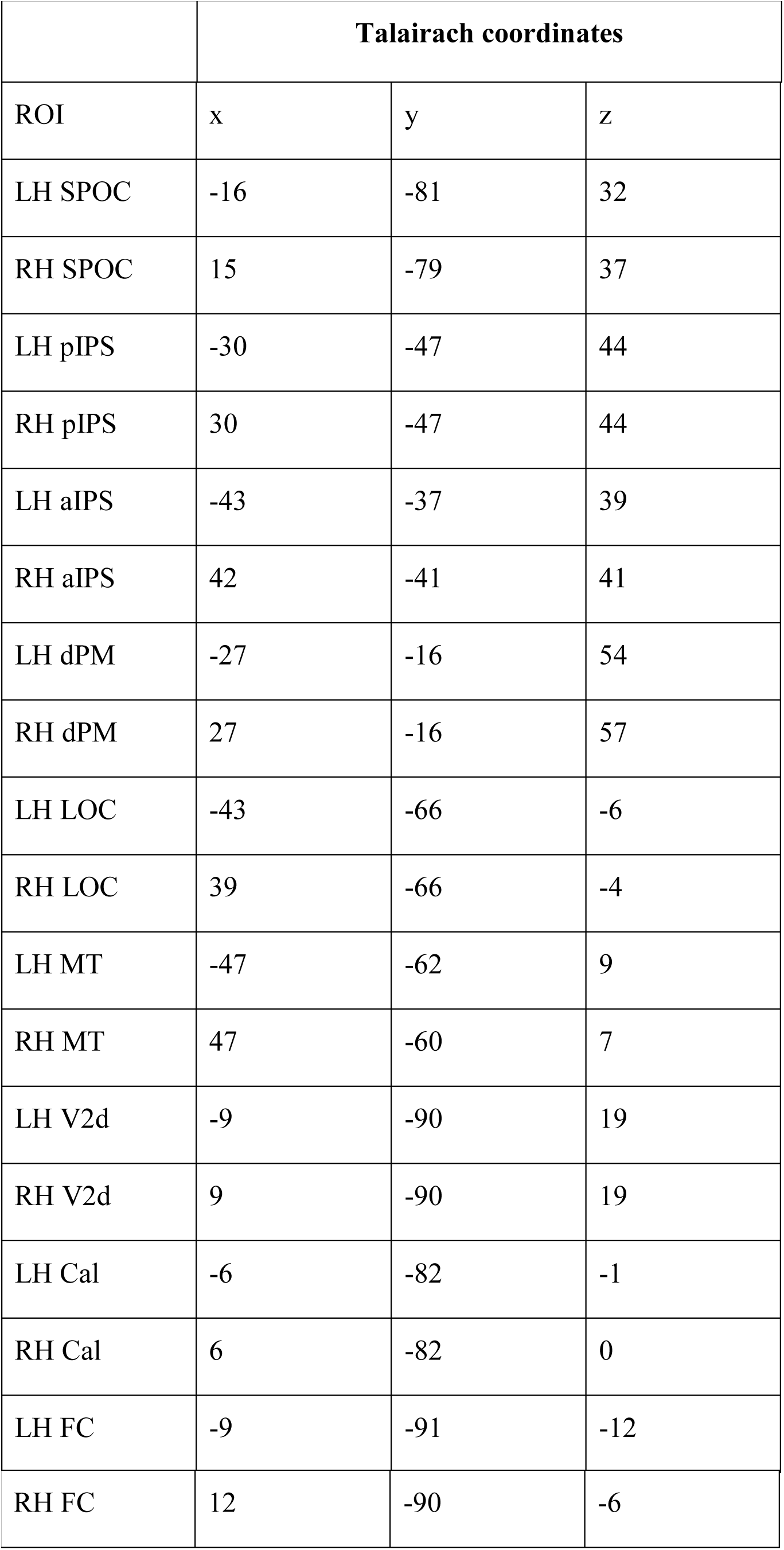
Average Talairach coordinates for each ROI.

|  | **Talairach coordinates** |  |  |
| --- | --- | --- | --- |
| ROI | x | y | z |
| LH SPOC | -16 | -81 | 32 |
| RH SPOC | 15 | -79 | 37 |
| LH pIPS | -30 | -47 | 44 |
| RH pIPS | 30 | -47 | 44 |
| LH aIPS | -43 | -37 | 39 |
| RH aIPS | 42 | -41 | 41 |
| LH dPM | -27 | -16 | 54 |
| RH dPM | 27 | -16 | 57 |
| LH LOC | -43 | -66 | -6 |
| RH LOC | 39 | -66 | -4 |
| LH MT | -47 | -62 | 9 |
| RH MT | 47 | -60 | 7 |
| LH V2d | -9 | -90 | 19 |
| RH V2d | 9 | -90 | 19 |
| LH Cal | -6 | -82 | -1 |
| RH Cal | 6 | -82 | 0 |
| LH FC | -9 | -91 | -12 |
| RH FC | 12 | -90 | -6 |

## Results

### Localizers and Univariate Results

The Talairach coordinates of each ROI are specified in Table 1. Activation maps for each voxelwise contrast used to localize our ROIs are shown in Figure 3. We used the univariate contrast (Action execution > baseline) to localize sensorimotor areas known to be part of the action network (Figure 3A). Retinotopic mapping allowed us to localize the parts of the visual cortex corresponding to the peripheral representations of the objects as well as central vision on a set of 14 participants (Figure 3B). While the Calcarine sulcus showed higher activation for eccentricities corresponding to the peripheral (6.7-10°) than central vision (1-4.5°), the Foveal cortex showed higher activation for eccentricities corresponding to central than peripheral vision. Area V2d was localized based on the openly available probabilistic atlas (Wang et al. 2015).

Before performing MVPA, we examined whether we could detect differences in processing objects features during the planning phase using the univariate contrast of: (Plan Align CCW-left > Plan Align CW-right) and (Plan Open reach CCW-left > Plan Open reach CW-right). These contrasts did not reveal any active voxel. This is expected given that participants were lying still and viewed both objects in the lower periphery simultaneously. Therefore, at this point of the task there was no difference in sensory or motor signals in the two conditions that could have elicited higher activation in one case than the other. Figure 2C shows a time course from the left calcarine which is representative of other EVC areas and areas of the action network. We then performed MVPA to examine differences in the pattern of activity.

### Decoding

Statistical values for each comparison are specified in Table 2. Means and standard deviations are indicated in Table 3. Figure 4 shows the percent decoding accuracy in each ROI for pairwise comparisons of object features (CCW-left vs. CW-right) during the planning and execution phase within Align and Open reach movements.

**Figure 4.**
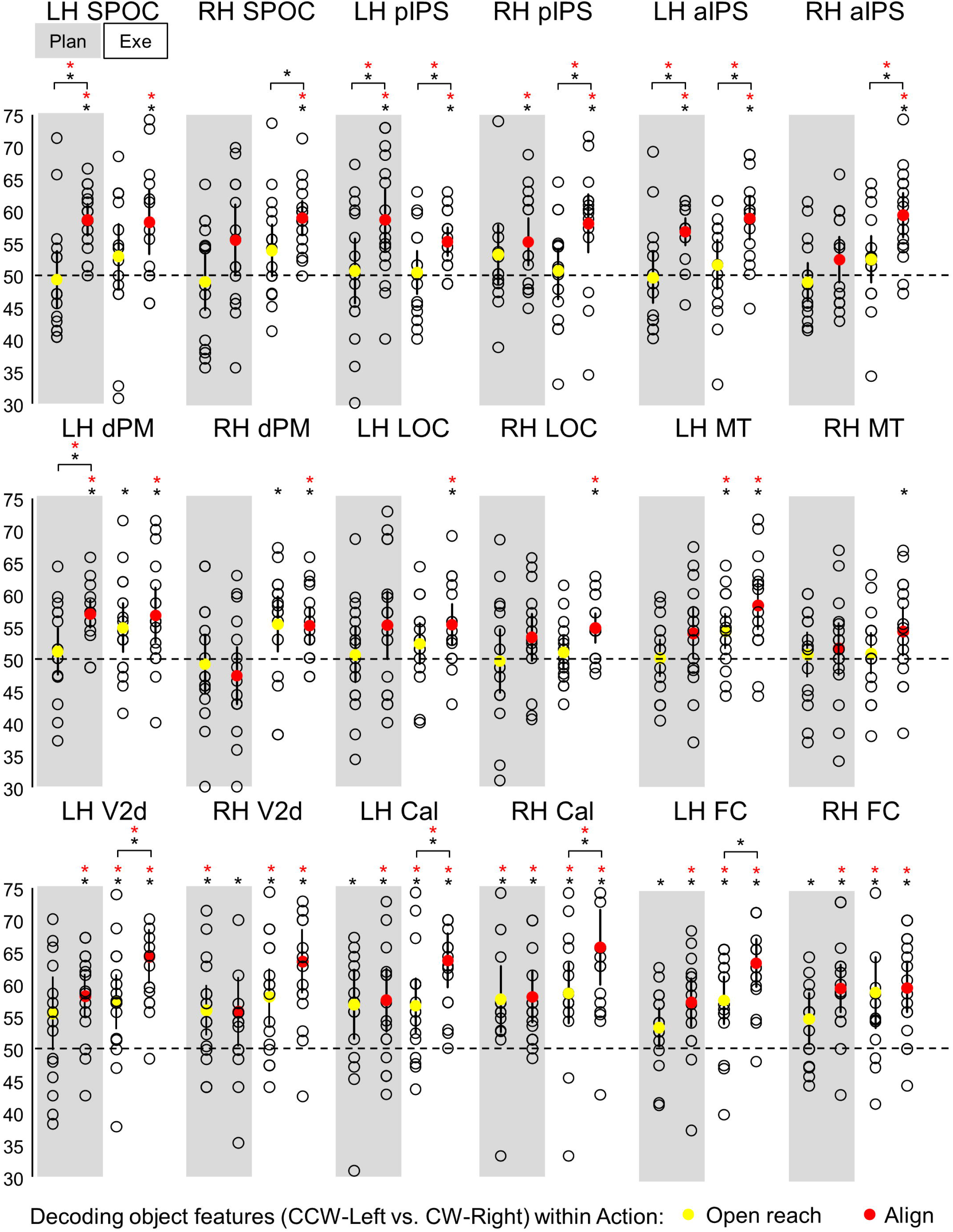
Classifier decoding accuracies for the discrimination between object features in our ROIs within Action type. The scatterplots show decoding accuracies for each participant along with the average (colored circles) for the dissociation between the two oriented objects (CCW-left vs. CW-right) during the planning (left cluster pair) and execution phase (right cluster pair) of Open reach trials (yellow circles) and Align trials (red circles). Chance level is indicated with a dashed line at 50% of decoding accuracy. Error bars show 95% confidence intervals. Black asterisks indicate statistical significance with two-tailed t-tests across subjects with respect to 50%. Red asterisks indicate statistical significance based on an FDR correction of q < .05.

**Table 2.**
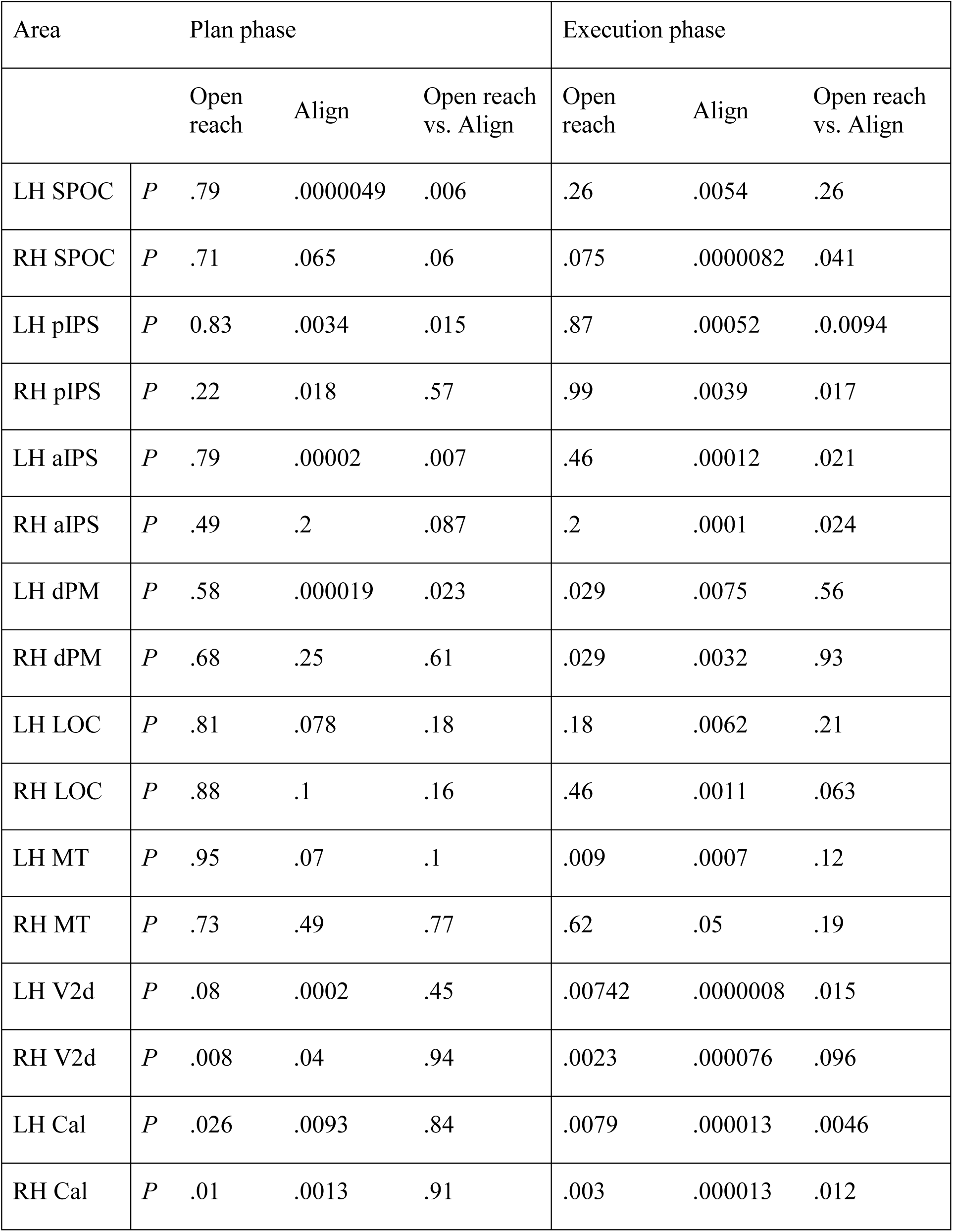
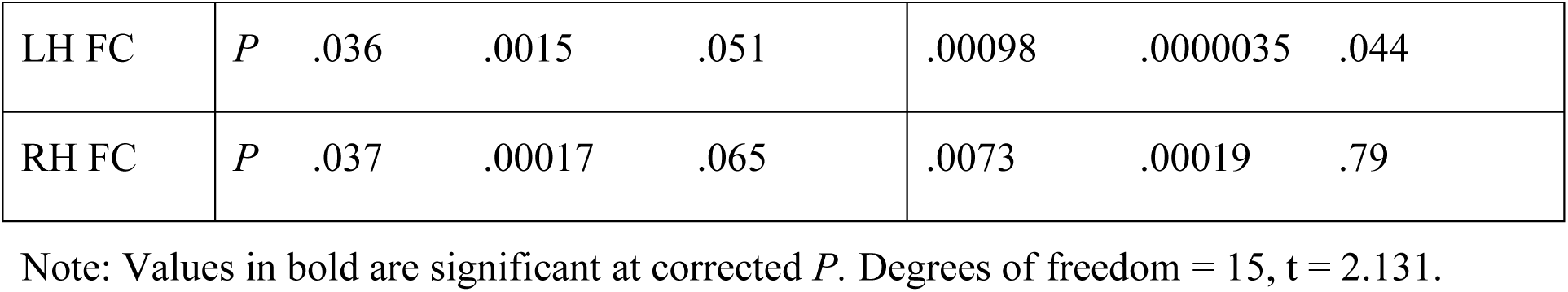
Statistical values for classifier decoding accuracies for the discrimination between object features (CCW-left vs. CW-right) against chance level.

| Area |  | Plan phase |  |  | Execution phase |  |  |
| --- | --- | --- | --- | --- | --- | --- | --- |
|  |  | Open reach | Align | Open reach vs. Align | Open reach | Align | Open reach vs. Align |
| LH SPOC | *P* | .79 | .0000049 | .006 | .26 | .0054 | .26 |
| RH SPOC | *P* | .71 | .065 | .06 | .075 | .0000082 | .041 |
| LH pIPS | *P* | 0.83 | .0034 | .015 | .87 | .00052 | .0.0094 |
| RH pIPS | *P* | .22 | .018 | .57 | .99 | .0039 | .017 |
| LH aIPS | *P* | .79 | .00002 | .007 | .46 | .00012 | .021 |
| RH aIPS | *P* | .49 | .2 | .087 | .2 | .0001 | .024 |
| LH dPM | *P* | .58 | .000019 | .023 | .029 | .0075 | .56 |
| RH dPM | *P* | .68 | .25 | .61 | .029 | .0032 | .93 |
| LH LOC | *P* | .81 | .078 | .18 | .18 | .0062 | .21 |
| RH LOC | *P* | .88 | .1 | .16 | .46 | .0011 | .063 |
| LH MT | *P* | .95 | .07 | .1 | .009 | .0007 | .12 |
| RH MT | *P* | .73 | .49 | .77 | .62 | .05 | .19 |
| LH V2d | *P* | .08 | .0002 | .45 | .00742 | .0000008 | .015 |
| RH V2d | *P* | .008 | .04 | .94 | .0023 | .000076 | .096 |
| LH Cal | *P* | .026 | .0093 | .84 | .0079 | .000013 | .0046 |
| RH Cal | *P* | .01 | .0013 | .91 | .003 | .000013 | .012 |
| LH FC | *P* | .036 | .0015 | .051 | .00098 | .0000035 | .044 |
| RH FC | *P* | .037 | .00017 | .065 | .0073 | .00019 | .79 |

Note: Values in bold are significant at corrected *P*. Degrees of freedom = 15, *t* = 2.131.

**Table 3.**
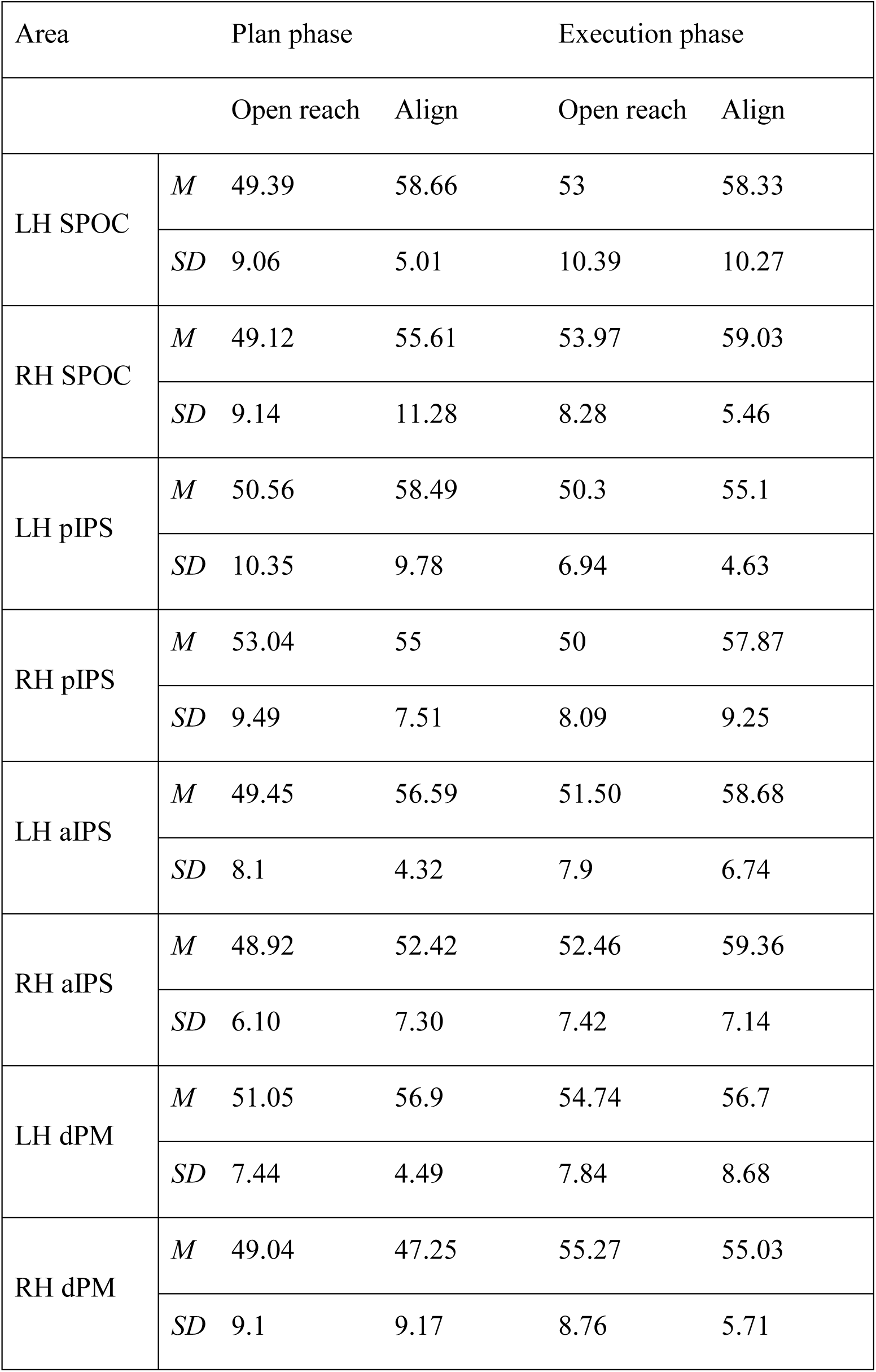
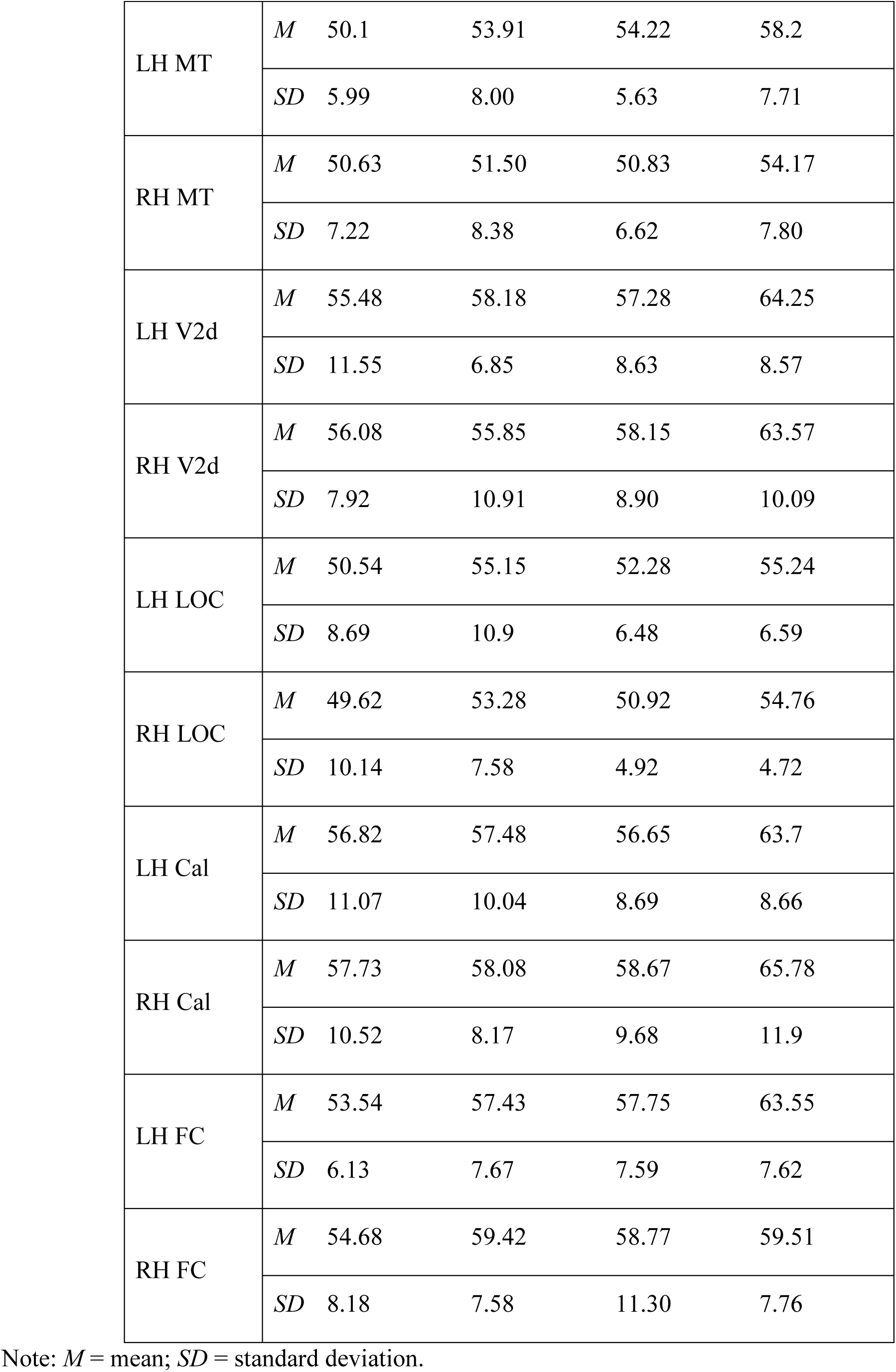
Means and standard deviations of decoding accuracies for the dissociation between object features (CCW-left vs. CW-right) for each condition in each ROI.

| Area |  | Plan phase |  | Execution phase |  |
| --- | --- | --- | --- | --- | --- |
|  |  | Open reach | Align | Open reach | Align |
| LH SPOC | *M* | 49.39 | 58.66 | 53 | 58.33 |
|  | *SD* | 9.06 | 5.01 | 10.39 | 10.27 |
| RH SPOC | *M* | 49.12 | 55.61 | 53.97 | 59.03 |
|  | *SD* | 9.14 | 11.28 | 8.28 | 5.46 |
| LH pIPS | *M* | 50.56 | 58.49 | 50.3 | 55.1 |
|  | *SD* | 10.35 | 9.78 | 6.94 | 4.63 |
| RH pIPS | *M* | 53.04 | 55 | 50 | 57.87 |
|  | *SD* | 9.49 | 7.51 | 8.09 | 9.25 |
| LH aIPS | *M* | 49.45 | 56.59 | 51.50 | 58.68 |
|  | *SD* | 8.1 | 4.32 | 7.9 | 6.74 |
| RH aIPS | *M* | 48.92 | 52.42 | 52.46 | 59.36 |
|  | *SD* | 6.10 | 7.30 | 7.42 | 7.14 |
| LH dPM | *M* | 51.05 | 56.9 | 54.74 | 56.7 |
|  | *SD* | 7.44 | 4.49 | 7.84 | 8.68 |
| RH dPM | *M* | 49.04 | 47.25 | 55.27 | 55.03 |
|  | *SD* | 9.1 | 9.17 | 8.76 | 5.71 |
| LH MT | *M* | 50.1 | 53.91 | 54.22 | 58.2 |
|  | *SD* | 5.99 | 8.00 | 5.63 | 7.71 |
| RH MT | *M* | 50.63 | 51.50 | 50.83 | 54.17 |
|  | *SD* | 7.22 | 8.38 | 6.62 | 7.80 |
| LH V2d | *M* | 55.48 | 58.18 | 57.28 | 64.25 |
|  | *SD* | 11.55 | 6.85 | 8.63 | 8.57 |
| RH V2d | *M* | 56.08 | 55.85 | 58.15 | 63.57 |
|  | *SD* | 7.92 | 10.91 | 8.90 | 10.09 |
| LH LOC | *M* | 50.54 | 55.15 | 52.28 | 55.24 |
|  | *SD* | 8.69 | 10.9 | 6.48 | 6.59 |
| RH LOC | *M* | 49.62 | 53.28 | 50.92 | 54.76 |
|  | *SD* | 10.14 | 7.58 | 4.92 | 4.72 |
| LH Cal | *M* | 56.82 | 57.48 | 56.65 | 63.7 |
|  | *SD* | 11.07 | 10.04 | 8.69 | 8.66 |
| RH Cal | *M* | 57.73 | 58.08 | 58.67 | 65.78 |
|  | *SD* | 10.52 | 8.17 | 9.68 | 11.9 |
| LH FC | *M* | 53.54 | 57.43 | 57.75 | 63.55 |
|  | *SD* | 6.13 | 7.67 | 7.59 | 7.62 |
| RH FC | *M* | 54.68 | 59.42 | 58.77 | 59.51 |
|  | *SD* | 8.18 | 7.58 | 11.30 | 7.76 |

Note: *M* = mean; *SD* = standard deviation.

As shown in Figure 4, during the planning phase, we could decode object orientation during the Align but not Open reach condition in bilateral pIPS and Foveal cortex, as well as in left SPOC, aIPS, dPM, V2d and Calcarine sulcus. However, only left SPOC, pIPS, aIPS and dPM area showed higher dissociation of object properties in Align than Open reach movements, suggesting an action-dependent representation of object features in these areas. In addition, the right Calcarine area showed significant dissociation of object features in Align and Open reach conditions, with no difference between movement types.

In the execution phase, we found significant decoding of object orientation for Align but not Open reach movements in bilateral SPOC, pIPS, aIPS, dPM and LOC. However, only bilateral pIPS and aIPS showed higher dissociation of object orientation in Align than Open reach movements. In addition, bilateral Calcarine area, Foveal cortex and left MT showed significant dissociation of object features in Align and Open reach conditions, as well as higher dissociation for Align than Open reach movements in bilateral Calcarine area, suggesting that vision of the moving hand enhanced dissociations in the EVC during moving execution.

### Cross-decoding

As shown in Figure 5, when the classifier was trained on Align and tested on Open reach trials, left V2d and Foveal cortex showed significant decoding accuracies in the plan phase, while right V2d showed significant decoding accuracies only in the execution phase. When the classifier was trained on Open reach trials and tested on Align trials, left V2d, Calcarine sulcus and Foveal cortex showed significant decoding accuracies in the execution phase.

**Figure 5.**
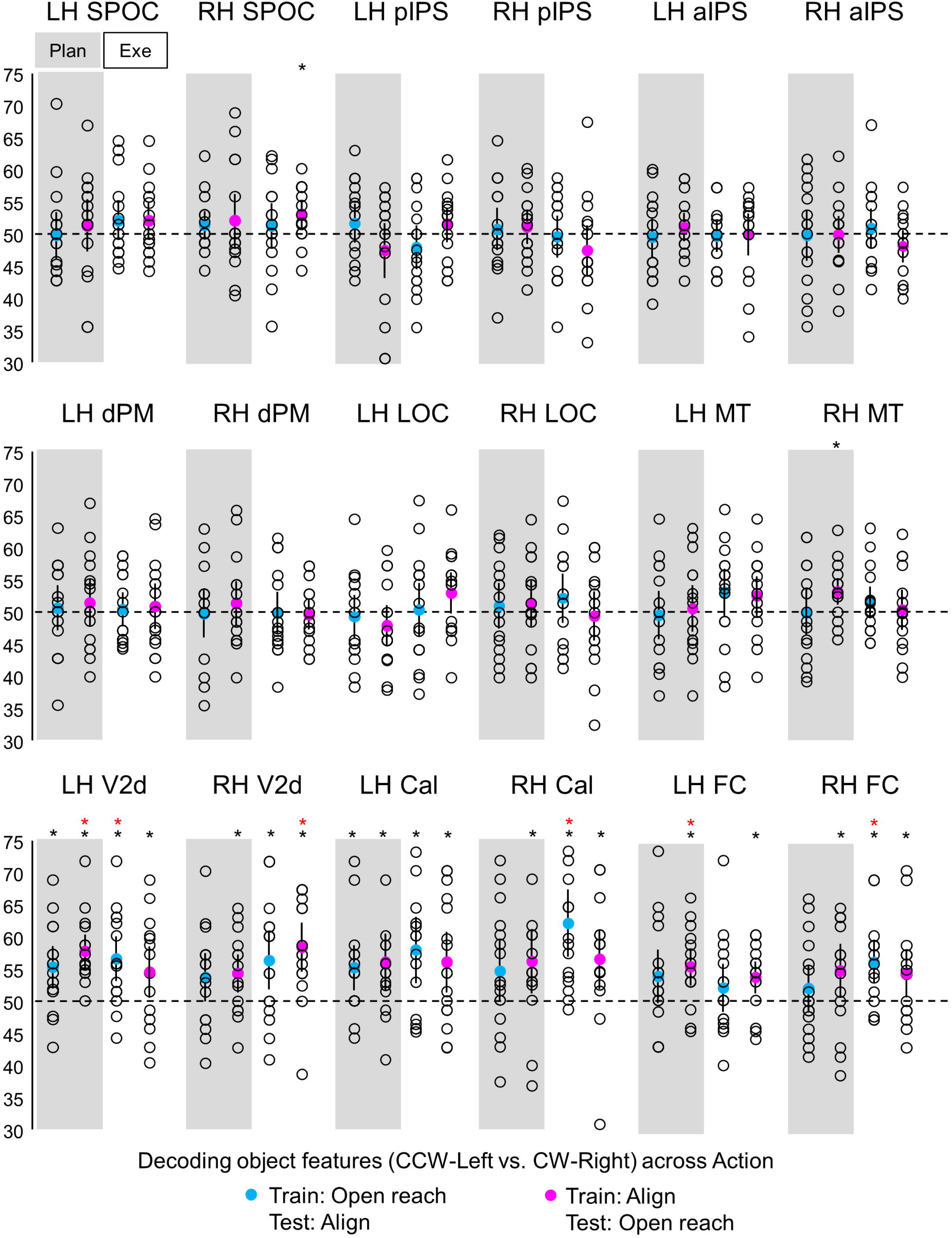
Classifier decoding accuracies for the discrimination between object features in our ROIs across Action type. The scatterplots show decoding accuracies for each participant along with the average (colored circles) for the dissociation between the two oriented objects (CCW-left vs. CW-right) during the planning (left cluster pair) and execution phase (right cluster pair). The blue circles indicate accuracies obtained when the classifier was trained on Open reach and tested on Align trials. The fuchsia circles indicate accuracies obtained when the classifier was trained on Align and tested on Open reach trials. Chance level is indicated with a dashed line at 50% of decoding accuracy. Error bars show 95% confidence intervals. Black asterisks indicate statistical significance with two-tailed t-tests across subjects with respect to 50%. Red asterisks indicate statistical significance based on an FDR correction of q < .05.

## Discussion

We examined whether and how early in the visual cortex the representation of object location and orientation is modulated by action planning. We found that during action planning and execution, the activity pattern in SPOC, pIPS, aIPS and dPM cortex allowed dissociating between the two concurrently presented objects in Align movements, which required orienting the hand according to object features, but not Open reach movements, which did not require adjusting hand orientation. Further, the dissociation was higher when planning an Align than an Open reach tasks. This suggests that upcoming actions enhance the representation of object properties that are relevant for the particular movement. Further, in the EVC we could reliably decode object features during the planning phase, but there was no difference between Align and Open reach movements. Taken together, these results provide a whole-brain, framework for understanding how action-relevant object features are cortically represented during action planning.

Our results show that the representation of objects features, such as orientation and location, is shaped by the upcoming action seconds before participants initiate a movement. This is likely due to the fact that different objects require different motor plans. Previous studies have shown the involvement of the fronto-parietal cortex in adjusting hand orientation during action execution. For instance, the superior end of the parietal occipital cortex in humans (SPOC) and macaques (V6A) has been shown to play a key role in processing wrist movements for hand orientation (Battaglini et al. 2002; Fattori et al. 2010; Monaco et al. 2011). Moreover, pIPS encodes 3D visual features of objects for hand actions in macaque (Sakata et al. 1998) and, along with dPM, it is involved in the adjustment of hand and wrist orientation during action execution in humans (Monaco et al. 2011). Although the aIPS is known to be highly involved in the pre-shaping of the fingers for grasping actions in humans (Culham et al. 2003; Cavina-Pratesi et al. 2010; Monaco et al. 2014, 2015, 2017) and macaques (for a review see (Turella and Lingnau 2014), neurophysiology studies indicated enhanced neuronal activity in this area when orienting the wrist during a grasping movement (Murata et al. 2000; Baumann et al. 2009). Our results extend previous findings by indicating that object orientation modulates the activity pattern in the fronto-parietal cortex not only during action execution, but also before initiating the action in a task-dependent manner.

Behavioral and neuroimaging research has revealed that the processing of action-relevant features can be enhanced during movement preparation, and that action planning can be decoded in the EVC (Bekkering and Neggers 2002; van Elk et al. 2010; Gutteling et al. 2015; Gallivan et al. 2019; Monaco et al. 2020). Therefore, we investigated whether object representation is influenced by the upcoming action as early as in the EVC during the planning phase preceding action execution. In addition to exploring the activity pattern in the Calcarine area corresponding to the retinotopic location of the objects below the fixation point, we also investigated the activity in the Foveal cortex, corresponding to central vision. In fact, the foveal cortex has been shown to contain visual information about objects presented in the visual periphery and this phenomenon has been found to correlate with task performance and to be critical for extra-foveal perception (Williams et al. 2008; Chambers et al. 2013; Fan et al. 2016). Our results show dissociable preparatory signals for different object orientations in the EVC. However, the decoding accuracy for the dissociation between object features did not differ between the two movement types (Align vs. Open reach), indicating that visuomotor planning toward oriented objects can be decoded in the EVC regardless of the type of action to be performed (Align or Open reach). During action execution there was higher decoding accuracy for the dissociation between object features in Align than Open reach tasks, however this result is most likely driven by the visual input of the moving hand in the participant’s visual field during the executing of the movement.

For areas that show a dissociation during the planning phase between the two orientations for Align as well as Open reach movements (i.e., the right Calcarine sulcus) we suggest that these areas predominantly receive location-relevant feedback. However, we cannot disambiguate whether object location or orientation drives the modulation, as the object on the left was oriented at about -45° while the one on the right was oriented approximately at 45°. In addition, both objects were present at the same time. As a consequence, our classification results may reflect tuning to object location, orientation, or both. However, in areas that show higher dissociation between the two objects for Align than Open Reach, such as SPOC, pIPS, aIPS and dPM, it is likely that object orientation allows for decoding, as it is relevant for Align movements but not coarse Open reach movements.

The cross-decoding analysis allowed determining whether object information is encoded in similar ways in the two Action types, or whether feature encoding is different across the two movements. Our results indicate that object features cannot be generalized across the two Action types in sensorimotor of dorsal and ventral visual stream. This is in line with the idea that the representation of object features is shaped by the upcoming action, even if the retinal inputs are exactly the same for different action plans. On the other hand, the results in left V2d and Foveal cortex showed that object representation can be generalized across action plans, but only when the classifier is trained on the Align and tested on the Open reach condition. This can be explained by the fact that object features encoding is enhanced in the Align condition, which requires detailed processing of object location and orientation, while the Reach condition only requires a coarse representation of the targets.

We filter feature information from redundant stimuli in the world around us to successfully plan actions. Behavioral and neural evidence has shown that attention is directed to the location of a planned movement (Bekkering and Neggers 2002; Moore et al. 2004). Since there is a tight linkage between attention and intention, researchers have suggested that these processes are subserved by the same neural mechanisms (Rizzolatti et al. 1987; Cattaneo and Rizzolatti 2009). Attentional mechanisms may in principle have contributed to our results in the EVC. However, the dissociation between object features was not limited to the peripheral visual field modulated by the stimulus (V1 and V2), but extended to the foveal cortex which was not directly stimulated. Further, attention would have likely driven stronger modulations for Align than Open reach tasks, as more effort and accurate processing was required for the former rather than the latter action. Yet, we found no difference in the modulations for Align and Open reach tasks in the EVC. An alternative interpretation of our results might be related to predictive remapping mechanisms by which some brain areas become responsive to the visual stimuli that will be brought into their receptive field by an upcoming saccade (Duhamel et al. 1992; Melcher 2007; Cavanagh et al. 2010). Although eye movements were not allowed during the experiment, it is possible that the natural tendency to look towards the direction of a stimulus might have initiated predictive remapping mechanisms. As such, the information processing might be related to an upcoming saccade planning. Further investigations are needed to determine the potential influence of saccade planning on the cortical representation of object features. Even though this fMRI setup did not allow recording eye movements, several behavioral studies have examined fixation when participants planned actions towards an object presented in the lower visual field. Participants, including naive ones, could reliably fixate on a point for long intervals during each phase of a trial (Gallivan, Mclean, et al. 2013; Monaco et al. 2020), suggesting that eye movements alone are unlikely to explain decoding results. The decoding results in area MT corroborate our suggestion by showing significantly different representations for Align and Open reach conditions in the execution phase, likely driven by the view of the moving hand, but not in the plan phase.

## Conclusion

To conclude, our results indicate that the representation of a workspace with two oriented targets varies as a function of the planned action. This is evident in dorsal stream areas, known to be specialized in action preparation, and to a lesser extent in the EVC. Our findings suggest a role of these areas in predictive coding of actions based on internal models that take into account visual and somatosensory anticipations of upcoming movements, as well as object features that are relevant for subsequent actions. This mechanism might be mediated by bidirectional functional connections between dorsal stream areas, known to be involved in action planning, and the EVC. Feedback connections from the dorsal stream areas might tune the signal in the EVC to enhance the representation of action-relevant object features.

## Acknowledgments

This project has received funding from the European Union’s Horizon 2020 research and innovation programme under the Marie Sklodowska-Curie grant agreement No 703597 to Simona Monaco.

## Abbreviations

SPOC: superior parietal occipital cortex
pIPS: posterior intraparietal sulcus
aIPS: anterior intraparietal sulcus
dPM: dorsal premotor area
LOC: lateral occipital complex
MT: middle temporal visual area
V2d: dorsal secondary visual cortex
Cal: Calcarine sulcus
FC: foveal cortex
LH: left hemisphere
RH: right hemisphere

## Conflict of Interest

The authors declare that the research was conducted in the absence of any commercial or financial relationships that could be construed as a potential conflict of interest.

## Author Contributions

SM conceived and designed the study. JVI and SM collected the fMRI data. JVI performed pre-processing, univariate analysis, and psychophysiological interaction analysis. SM performed multivariate analysis. JVI wrote the first draft of the manuscript. SM wrote sections of the manuscript and revised it. All authors read, revised, and approved the submitted version of the manuscript.

## Data availability Statement

The data that support the findings of this study are available from the corresponding author upon request.

## Notes

### Competing Interest Statement

The authors have declared no competing interest.

